# Reduction of *Listeria* on stainless steel surfaces is impacted by sanitizer application method

**DOI:** 10.1101/2025.04.09.647964

**Authors:** Yang Jiao, Jakob Baker, Calvin Slaughter, Devin Daeschel, Abigail B. Snyder

**Author notes:** Address correspondence to Abigail B. Snyder.

## Abstract

Pathogen cross-contamination during food production is controlled through sanitation. However, sanitizer efficacy is often studied in bench-scale experiments (e.g., submerged coupons in static or stirred sanitizer) which poorly approximate fluid dynamics. This limits our understanding of how effective sanitization is in commercial application. This study paired computational fluid dynamic (CFD) estimates of shear stress during spray application of sanitizer with measurements of *Listeria innocua* reduction on stainless steel by 100 ppm hypochlorite sanitizer under various application methods. Static submersion of inoculated coupons for 3 s resulted in a log reduction of 2.3 ± 0.1 log CFU. Bench-scale spray application for 3 s had the largest microbial reduction at the point of sanitizer spray impingement (7.5 ± 0.5 log CFU) and directly adjacent to the impingement point (6.4 ± 0.7 log CFU) where shear stress was the highest. Surface locations below the impingement point that only received fluid film sanitizer run-off had a significantly lower microbial reduction of 0.4 ± 0.1 log CFU (p < 0.05). At the pilot scale, sanitizer spray manually applied by operators achieved a 2.5 ± 0.4 log CFU reduction, which was significantly lower than what was achieved during bench-scale spray application (p < 0.05). Microbial reduction from manual operation of spray equipment was also significantly different among operators (p < 0.05). Discrepancies between bench-scale spraying, pilot-scale spraying, and submerged coupons underscores the need for sanitizer validation under realistic conditions to better understand the risk reduction achieved through sanitation programs during food processing.

## 1. INTRODUCTION

Cross-contamination of pathogens from environmental surfaces to food during processing has contributed to numerous recalls and foodborne illness outbreaks (FDA, 2025; Nüesch□Inderbinen et al., 2021; USDA, 2025). Pathogens like *Listeria monocytogenes* can persist on environmental surfaces despite sanitation operations and increase the risk of product cross-contamination (Daeschel et al., 2022; Elson et al., 2019; Ferreira et al., 2014; Nüesch□Inderbinen et al., 2021; Todd & Notermans, 2011). Environmental surface sanitation is the primary control against cross-contamination of food from *Listeria monocytogenes* and other environmental pathogens. Environmental surfaces in food manufacturing plants include the food-contact and non-food-contact surfaces of (1) processing equipment and utensils, (2) non-processing equipment and tools such as push carts and floor mats, as well as (3) the building’s structures such as floors and drains (Feng et al., 2023). Apart from the internal surfaces of equipment that can be subjected to clean-in-place systems, environmental surfaces are typically manually cleaned and sanitized. Cleaning techniques include wiping, scraping, brushing, spraying, and washing of surfaces to physically remove food residue and microbial populations from surfaces. Sanitizing is then often accomplished by spraying surfaces with a sanitizer to chemically inactivate the remaining microbial populations.

The efficacy of surface sanitization through sanitizer spraying is dependent on physical treatment variables such as the contact time, shear stress, and mode of application (Al Saabi et al., 2021; Cai et al., 2020; Goode et al., 2018). In practice, sanitizing surfaces is a dynamic process with variable shear stress and contact time. Some surface locations receive the full force from the sanitizer spray directly at the point of surface impingement. Locations adjacent to the impingement point are exposed to sanitizer that splashes or flows outward with diminished force. Other locations are exposed to sanitizer that flows down from the point of impingement*’* due to gravity, providing even lower shear forces on the surface. Furthermore, since sanitizer sprayers are manually operated by personnel, the application of the treatment varies depending on individual techniques, including differences in duration, nozzle proximity to surface, and consistency at each location on the surface. Prior bench-scale research on the efficacy of sanitizers has often neglected the role of shear stress on microbial reduction, opting for inoculated surface immersion within sanitizer solutions (Kim et al., 2023; Ruiz□Llacsahuanga et al., 2022) or immersions of pure culture within test tubes of sanitizer (OCSPP, 2012). Similarly, pilot-scale challenge studies that have manually applied sanitizer sprays to surfaces fail to quantify the different shear stress levels relevant to the range of exposure scenarios at different locations across a surface (Bailey et al., 1986; X. Wang et al., 2024). Therefore, there is a gap in the research on environmental sanitation regarding the impact of shear stress from sanitizer sprays and how this translates to risk reduction. The goal of this study was to quantify the shear stress and microbial reduction at different locations on a stainless steel surface with respect to the point of sanitizer spray impingement. In addition, we performed a pilot-scale challenge study to quantify the observed reduction in *Listeria* on a surface when sprayers were manually operated by researchers.

## 2. Materials and methods

### 2.1 Bench-scale spray-chamber construction

A sprayer (#FI-5N, Foamit, Grand Rapids, MI, USA) was connected to a 19 L (5 gallon) reservoir with a 5 m hose. The sprayer was also connected to a compressed air system with 50 psi pressure, which generated a volume flow rate of ∼ 2 L/min. The nozzle of the sprayer was a flat-fan shape VeeJet nozzle (H1/4U-SS2520). A bench-scale sanitation chamber (38 × 38 × 38 cm^3^) was custom built for sanitizer spray application onto a vertically fixed 30 × 30 cm^2^ surface.

The frame was made with Polyvinyl Chloride (PVC) tubes and wrapped with PVC plastic sheets to enclose the spray from splashing to the surrounding environment. The enclosure had an opening at the front and a PVC tube was attached horizontally approximately 24 cm from the frame’s bottom to secure the sprayer’s position, ensuring precise control and consistent replicates during the spraying experiments. A manufactured PVC coupon (30 × 30 cm^2^) was tied at the back of the frame to secure interchangeable stainless steel coupons (30 × 30 cm^2^) at a distance of 38 cm directly in front of the sprayer nozzle.

### 2.2 Computational fluid dynamics modeling

A computational fluid dynamics model (CFD) was constructed in ANSYS Fluent (ANSYS, 2022) to quantify shear stress at different locations on a surface relative to the point of sanitizer spray impingement. Validation was done with experimental flow pattern and velocity field. The geometry was plotted based on the exact components and dimensions of the bench-scale experimental setup as described in *section 2.1* to simulate the sanitizer spray and flow from the nozzle tip to the stainless steel surface. A control-volume approach was employed to integrate the governing equation over each cell in the mesh, and to allow for a set of discrete algebraic equations to be solved for the flow variables, including the pressure, velocity and shear stress.

#### 2.2.1 Governing equations and simplifications in geometry

To avoid complex geometry creation and many mesh grids in simulation, the sanitizer flow was simplified to a high-speed sanitizer column exiting from a custom-shaped pressurized outlet and impinging to the stainless steel wall. Volume of fraction (VOF) module was selected in the simulation since the scenario was simplified as liquid (sanitizer) entering a gas zone (air).

For the high-speed injection regarding turbulent flow, a *k-ε* turbulence model was selected for solving the constructed model, with specific equations as follows:

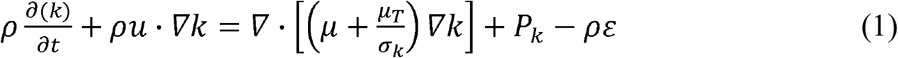

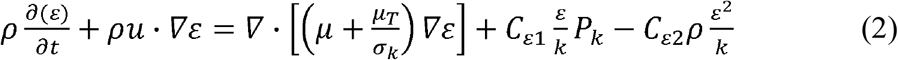

The standard *k-ε* turbulence model was solved using a pressure-based, double-precision implicit solver with second-order discretization for pressure and momentum. The convergence of the simulation was monitored by the continuity residual, residual of the turbulent kinetic energy (*k*), and residual of the turbulent dissipation (*ε*). Convergence was determined based on a calculated residual less than the tolerance of 1 × 10^−4^.

#### 2.2.2 Mesh generation

Fluent generated poly-hexacore mesh was applied to the whole domain. The mesh size was refined until the solved liquid velocity has reached a difference below 0.5% for finalization. The maximum and minimum size of cell was 0.003 and 0.002 m, respectively. At the pressure inlet, the mesh was refined to a maximum size of 0.002 m and a minimum size of 0.00025 m for converging. A 4-layer boundary layer was set at the wall with the excess ratio set to 0.272 and the growth rate set to 1.2. The computed dimensionless wall distance (*y*^*+*^) was less than 5 and confirmed that the designed mesh size was appropriate to apply the standard *k-ε* turbulence model for solving the wall shear stress (S. J. Wang & Mujumdar, 2005). The total number of cells generated in the whole domain was estimated as 2,881,612.

#### 2.2.3 Initial and boundary conditions

The flow boundary condition at the outlet of the nozzle was set as “pressure inlet” with a gauge pressure of 50 psi (3.45×10^5^ Pa) according to the experimental setup. The nozzle surface and stainless steel coupon surface were set as “wall,” and all other boundaries were set as “pressure outlet” with a gauge pressure of *P* = 0. Non-slip boundary condition was applied, and the properties of the sanitizer solution were set as the built-in definition of liquid water in the software since the dilute solutions have similar flow properties as water at room temperature (Fan et al., 2021) (Table 1). In the VOF model, the initial content of the spraying chamber was set as “gas” phase (*v* = 1), and the liquid column within the nozzle was set as “liquid” phase (*v* = 0).

**Table 1.** Fluid properties of air and liquid sanitizer in CFD modeling (25°C)

| Parameters | Species | Value |
| --- | --- | --- |
| Viscosity | Air | $10^{-6}$ Pa·s |
| | Sanitizer (100 ppm) | $10^{-3}$ Pa·s |
| Density | Air | $1.003 \text{ kg/m}^3$ |
| | Sanitizer (100 ppm) | $998 \text{ kg/m}^3$ |
| Contact angle | Between air and liquid sanitizer | $0.072 \text{ N}\cdot\text{m}$ |

#### 2.2.4 Post-processing

After convergence, the velocity of sanitizer liquid distribution and the shear stress on the coupon were analyzed directly from ANSYS. Velocity profiles (*u*) of sanitizer were obtained from the top and side view over the chamber as a 2-D contour plot, and the shear stress (τ_*w*_) over *x-* and *y-*direction on the coupon surface were obtained and explored as a 1-D line plot.

#### 2.2.5 Model validation

A high-speed camera (FASTCAM, Photron, San Diego, USA) was utilized to capture the velocity of sanitizer flow during treatment, and data was used to verify the effectiveness of the CFD model. The top view of the flow was captured consecutively, and the image was analyzed with “ImageJ” software (NIH image, USA) for estimating the flow velocity and to compare to the simulation results. The spray angle and spray width on the coupon surface were also obtained from both experiment and simulation, and alignment between those results also supported model validation.

### 2.3 Surface preparation

Stainless steel sheets (0.46 mm, 316/316 L, cold roll 2B) were cut into coupons with dimensions of 30 × 30 cm^2^, and 10 × 10 cm^2^ by the Machine Shop at Cornell University. The coupons were sterilized in an autoclave (Steris®, Mentor, OH, USA) at 121°C for 30 min under 15 psi. For pilot-scale studies, a stainless steel table with a top surface with a *length* × *width* of 120 × 70 cm^2^ was set on its side perpendicular to the floor. The stainless steel table was uniformly sprayed with ethanol (70% EtOH) and wiped with paper towels, then air-dried for 10 min.

### 2.4 Sanitizer preparation

Sanitizer solutions with 100 ppm of available chlorine were prepared using the XY-12 (Ecolab, St. Paul, CA, USA), a chlorine based liquid sanitizer with 8.4% sodium hypochlorite. The concentration was confirmed with a chlorine test kit (FryOilSaver Co., Matthews, NC, USA). This product and the concentration of 100 ppm sodium hypochlorite sanitizer was selected based on its common use in food processing environments (Mazaheri et al., 2023).

### 2.5 Inoculum preparation

Three strains (ATCC 51742, FSL C2-008, FSL A5-0424) of *Listeria innocua* were selected for this study from the Food Safety Laboratory at Cornell University (Ithaca, NY) based on prior studies with surface detachment (Chen & Snyder, 2023; Hsu et al., 2013; Pietrysiak & Ganjyal, 2018). Each strain was stored at -80°C in 15% glycerol (v/v). A loopful (10 µL) of frozen stock for each strain was cultured in brain heart infusion (BHI) broth (BD, Thermo Fisher Scientific, Waltham, MA, USA) by incubation at 37±2°C for 24±3 h with three successive transfers (24 h intervals), then streaked onto BHI agar plates and incubated at 37±2°C for 24 h. An isolated colony from each plate was inoculated into 2 × 5 mL BHI broth and incubated at 37 ± 2°C for 24 h. The culture was centrifuged at 5,000 RPM for 10 min (Eppendorf 5804R, Eppendorf, NY, USA), and the pellet was washed twice with 0.1% phosphate buffered saline (PBS) (BD, Thermo Fisher Scientific, Waltham, MA, USA). After washing, the cell pellets were resuspended in 10% volume of PBS to achieve a cell concentration of ∼8-log CFU/mL. A cocktail was prepared by mixing equal volumes of the three cultures.

### 2.6 Dried-down spot inoculation of surfaces

#### 2.6.1 Inoculation of coupons for submersion experiments

Cocktails were spot inoculated (0.1 mL) onto the center of stainless steel coupons (10 × 10 cm^2^). Then, the inoculated coupon was placed in a biosafety hood (Thermo Fisher Scientific, Waltham, MA, USA) (22°C) to dry down for 3 h prior to treatment. All experiments were conducted in biological triplicate.

#### 2.6.2 Inoculation of surfaces for bench-scale spray chamber experiments

Three locations on a stainless steel coupon (30 × 30 cm^2^) were selected as inoculation sites for the bench-scale sanitizer spraying experiment. The locations were named as “impingement,” “impingement-adjacent,” and “fluid-film,” indicating their position relative to the sanitizer spray’s impingement point on the surface. During spraying, the “impingement” location was at the center (0 cm) of the spray impingement on the surface, the “impingement-adjacent” position was 5.1 cm (2 in) to the left of the “impingement” point. The “fluid-film” location was 7.6 cm (3 in) below the “impingement” location. The surface was spot inoculated (0.1 mL) at each location and then the inoculated surface was placed in a biosafety hood (Thermo Fisher Scientific, Waltham, MA, USA) (22°C) to dry down for 3 h prior to treatment. All treatments were conducted in biological triplicate.

#### 2.6.3 Pilot-scale surface inoculation

The *L. innocua* cocktail was inoculated (10 μL spot × 10 spots) on the top of the stainless steel tables while they were in an upright position. The spot locations were predetermined and consistent among replicates. Inoculated surfaces were allowed to air dry at 22°C for 3 h while upright, then placed perpendicular to the floor for treatment. All treatments were conducted in biological triplicate.

### 2.7 Sanitizer treatments

#### 2.7.1 Treatment of surfaces by submersion into static sanitizer

Coupons were submerged for 3 s or 120 s of application time in 1 L of prepared 100 ppm sodium hypochlorite sanitizer. Following treatments that did not include a subsequent hold period for extended contact time after removal from the sanitizer, coupons were immediately processed according to the steps in *2*.*8 Recovery and enumeration*. Following treatments with subsequent hold periods to extend contact times, coupons were stored vertically for 120 s before being processed according to the steps in *2*.*8 Recovery and enumeration*

#### 2.7.2 Treatment of surfaces in a bench-scale spray-chamber

A bench-scale study was conducted by spraying the prepared sanitizer from a fixed position perpendicularly at vertically-fixed surfaces. This experiment assesses microbial reduction from the impinging fluid on the surface at 3 discrete locations (Impingement, Impingement-adjacent, and Fluid-film). The sprayer was operated for 3 s at a distance 38 cm from the surface. Additional experiments were conducted varying the treatment by: (1) prolonging spray application from 3 s to 2 min, as well as (2) maintaining a 3 s spray but extended the contact time to 2 min before swabbing and neutralizing.

#### 2.7.3 Treatment of surfaces by pilot-scale, manual spray application

Three researchers were instructed to sanitize the surface as would sanitation personnel without knowledge of the inoculation sites and assuming this scenario was within a food processing environment. The researchers were informed to tie back all hair, wear long pants, closed-toed shoes (not cloth shoes), as well as don splash-proof lab goggles, nitrile gloves, and a lab coat. No additional instructions were provided besides the safety protocol for manual operation of the sprayer. The distance (cm) between the sprayer nozzle and surface, the duration (s) of spraying treatment, and the application path of the sanitizer spray on the inoculated surface were extracted from video recordings of the treatments using an Apple iPhone XR (Apple Inc., Cupertino, CA, USA). Each researcher determined their own spray path, treatment duration, and nozzle distance from the surface; however, these variables were approximately the same for each researcher among their three replicate treatments.

### 2.8 Recovery and enumeration

Surfaces were swabbed with a sponge-stick with 10 mL Dey/Engley (D/E) neutralizing broth (SSL10DE, Neogen, St Paul, USA). Swabbing was conducted both vertically and horizontally (5 top to bottom strokes, followed by 5 left to right strokes). This swabbing protocol was validated in preliminary studies to ensure consistently accurate microbial counts. The swabs were stomached for 2 min at 260 rpm (Stomacher® 400 Lab Blender Series, Seward, UK). Tenfold serial dilutions were conducted, then 100 μL was spread onto BHI plates and incubated at 37°C for 24 h.

### 2.9 Statistical analysis

All statistical analyses were conducted in RStudio (R version 4.4.3). A one-way ANOVA, followed by Tukey’s post-hoc analysis, was used to compare the log reductions achieved by the different researchers during the pilot-scale sanitizing process. One-way Welch’s ANOVA, followed by Games-Howell post-hoc analysis, was performed to compare log reductions between the “Impingement,” “Impingement-adjacent,” and “Fluid-film” locations, as well as to assess the effects of different application and contact time parameters. Welch’s ANOVA and Games-Howell post-hoc analysis were also used to compare static sanitizer application with the spray application methodologies. Welch’s ANOVA and Games-Howell post-hoc analysis were chosen for these comparisons due to significant differences in the magnitude of variances among the groups of interest which violated the homogeneity of variance assumption required for standard one-way ANOVA and Tukey’s post-hoc analysis (Lee & Lee, 2018).

## 3. Results and Discussion

### 3.1 Shear stress varied based on location relative to the point of impingement during spraying

Shear stress estimation at the “Impingement,” “Impingement-adjacent,” and “Fluid-film” locations via CFD are shown in Figure 1B and Figure 1C. Shear stress was greatest at the “Impingement-adjacent” position at 75 Pa, followed by the “Impingement” location at 25 Pa, and lastly the ‘fluid-film’ position at 4 Pa. Although the force of the spray was greatest at the point of impingement, shear stress was greater directly adjacent to the impingement point. This is typical of shear stress measurements around the point of impingement in similar systems (Tu & Wood, 1996) due to the reduced velocity of the gas or liquid directly at the point of the impingement on the surface. Velocity then peaks directly adjacent to the impingement point and then reduces further at the “Fluid-film” location where we identified the lowest shear stress.

**Figure 1.**
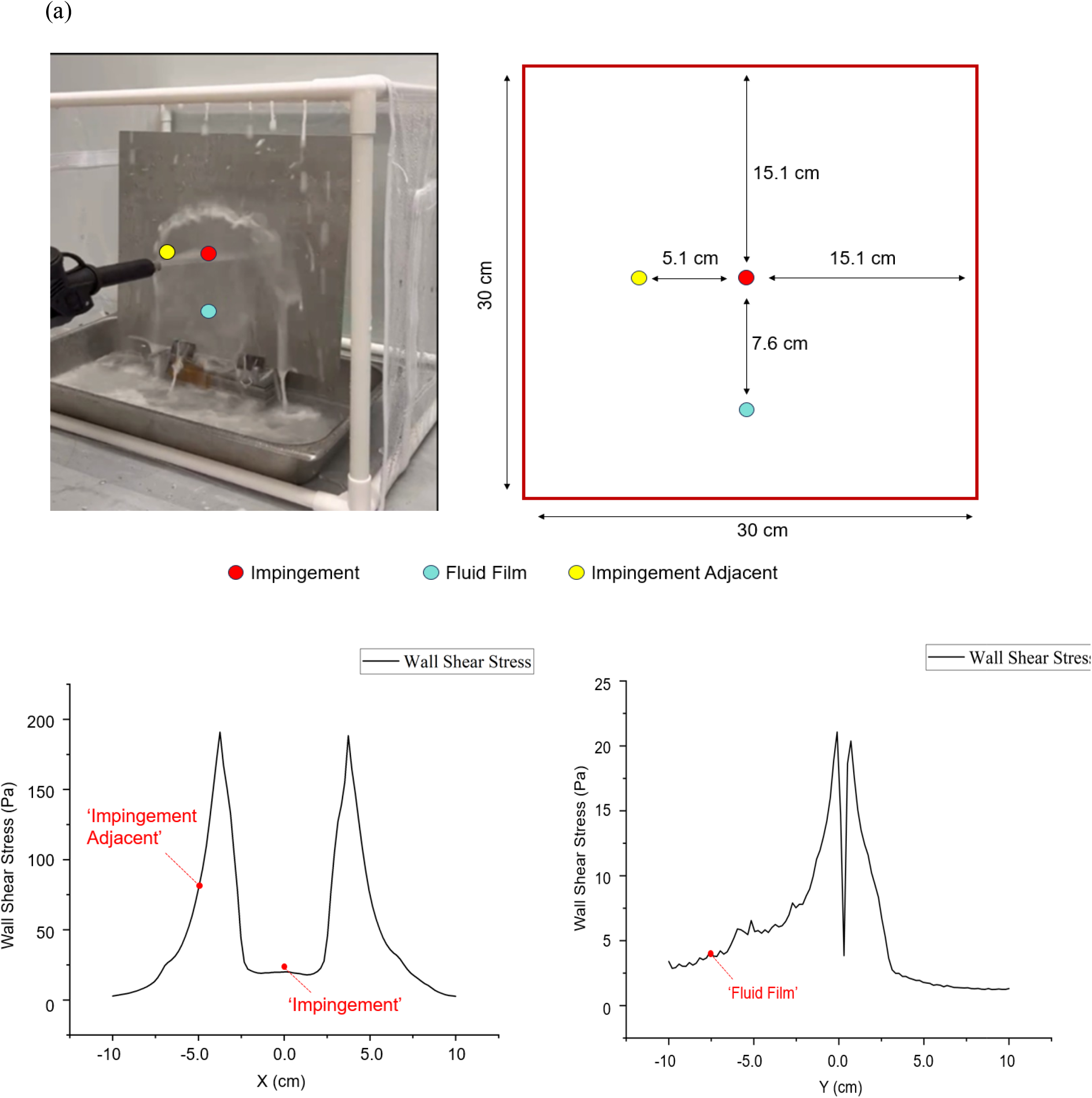
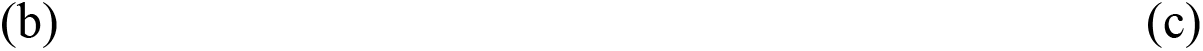
(a) Benchtop spray chamber configuration with inoculation sites. CFD simulation results for the (b) top view of the shear stress profile and (c) side view of shear stress profile.

### 3.2 Bacterial reduction from spraying was greatest at, and adjacent to, the point of impingement for both water and sanitizer sprays

During rapid exposure (3 s application time – 0 s extended contact time after cessation of spraying), average microbial reduction via a commercial sprayer in the benchtop spray chamber was the greatest at the “Impingement” location followed by the “Impingement-adjacent” location and lowest at the “Fluid-film” position for both the water and sanitizer treatments (Table 2). The log reduction from spray application of water at the “Impingement” position was 3.4 ± 0.6 log CFU/surface, at the “Impingement-adjacent” position was 3.2 ± 0.5 log CFU/surface, and at the “Fluid-film” position was 0.4 ± 0.8 log CFU/surface. These differences were not statistically significant (p > 0.05; Welch’s ANOVA; Games-Howell). For surfaces treated by the sprayer with 100 ppm of pressurized sanitizer, the log-reduction was again highest at the “Impingement” location followed the “Impingement-adjacent” and “Fluid-film” locations at 7.5 ± 0.5 log CFU/surface, 6.4 ± 0.7 log CFU/surface, and 0.4±0.1 log CFU/surface, respectively. Welch’s ANOVA followed by Games-Howell post-hoc analysis indicated a significant difference in microbial reduction between the “Impingement” and “Fluid-film” locations during sanitizer treatment. A coupon submerged in static sanitizer with no shear forces had 2.3 ± 0.1 log-reduction, which was greater than the “Fluid-film” location (p < 0.05) but lower than the “Impingement-adjacent” and “Impingement” locations (*p* < 0.05). Microbial reduction increased with the use of sanitizer compared to water during spraying for all locations on the surface (p < 0.05), but water spraying alone still managed to achieve a 3-log reduction on surfaces.

**Table 2.** Shear stress and log-reductions of *Listeria innocua* from submerged coupon and benchtop-spray chamber trials at 3 s application time, 0 s contact time. Values in a column not connected by the same letter are statistically different from one another (p < 0.05; Welch’s-ANOVA; Games-Howell)

| Site | Treatment | Shear Stress (Pa) | Log-Reduction (SE) |
| --- | --- | --- | --- |
| Submerged coupon | Sanitizer | 0 | 2.3 (0.1)<sup>AB</sup> |
| Impingement | Sanitizer | 25 | 7.5 (0.5)<sup>C</sup> |
| Impingement-adjacent | Sanitizer | 75 | 6.4 (0.7)<sup>AC</sup> |
| Fluid-film | Sanitizer | 4 | 0.4 (0.1)<sup>D</sup> |
| Impingement | Water | 25 | 3.4 (0.6)<sup>AB</sup> |
| Impingement-adjacent | Water | 75 | 3.2 (0.5)<sup>ABD</sup> |
| Fluid-film | Water | 4 | 0.4 (0.8)<sup>BD</sup> |

Unexpectedly, microbial reduction was highest (although not significantly different) at the “Impingement” location despite higher modeled shear stresses at the “Impingement-adjacent” location. This could be due to larger normal forces from the sanitizer spray directly at the point of impingement. It is also possible that the simulation model, which simulated shear stress under ideal flow conditions, underestimated the shear stress at the site of impingement due to deviations from CFD predicted flow that occur in the physical world (Grace & Taghipour, 2003). Alternatively, it may be the case that microbial reduction peaks after a certain level of shear force, which would explain why there was not a significant difference in microbial reduction between the “Impingement” and “Impingement-adjacent” locations. Indeed, Gibson et al. (1999) reported approximately a 3 log CFU/cm^2^ and 2 log CFU/cm^2^ reduction of surface-adhered *Staphylococcus aureus* and *Pseudomonas aeruginosa* biofilms, respectively, from a stainless steel surface with a pressurized water sprayer system and did not see this value increase after increasing the pressure of the sprayer from 17 to 70 bar. As expected, the “Fluid-film” location had the lowest microbial reduction during spray application likely due to the minimal shear forces at this location. Prior studies confirm that fluid films on surfaces from falling impinging streams generate less wall shear stress, limiting the mechanical reduction of bacterial populations attached to surfaces (Cai et al., 2020)) However, higher microbial reduction was observed for the coupon immersed in sanitizer despite having less shear stress than the “Fluid-film” location. Therefore, there may be differences other than shear stress between submersion and spray application of sanitizer that affect microbial reduction such as different rates of sanitizer degradation due to variable temperature or light exposure (Marriott & Gravani, 2006) or greater release of active ingredients during sanitizer impingement.

### 3.3 Microbial reduction at the “Fluid-film” location increased with sanitizer application time and contact time

The microbial reduction at the “Fluid-film” location increased from 0.4±0.1 to 2.0 ± 0.1 CFU log-reduction when a 120 s contact time was added following the cessation of the 3 s spray (Table 3). Increasing the spray application time to 120 s further increased the log-reduction per surface at the “Fluid-film” location to 6.3±0.3 CFU log-reduction per surface (*p* < 0.05) (Table 3). Submersion of a coupon in static sanitizer yielded higher log-reductions than was obtained at the “Fluid-film” location following the 3 s application + 0 s contact time and the 3 s + 120 s contact time treatments.

**Table 3.** *Listeria innocua* reduction on submerged coupons and at the “Fluid-film location under varying sanitizer application and contact times. Values in a column denoted with different letters are statistically different from one another (p < 0.05; Welch’s-ANOVA; Games-Howell)

| Site | Application Time (s) | Contact Time (s) | Log-Reduction (SE) |
| --- | --- | --- | --- |
| Submerged coupon | 3 | 0 | 2.3 (0.1)<sup>A</sup> |
|  | 3 | 120 | 4.4 (0.6)<sup>ABC</sup> |
|  | 120 | 0 | 6.0 (< 0.1)<sup>CD</sup> |
| Fluid-film | 3 | 0 | 0.4 (0.1)<sup>B</sup> |
|  | 3 | 120 | 2.0 (0.1)<sup>AC</sup> |
|  | 120 | 0 | 6.3 (0.3)<sup>D</sup> |

Although the literature is limited regarding the application of sanitizer sprayers on surfaces, some research has been done exploring the role of contact time (Falcó et al., 2018; Gélinas et al., 1984). However, the role of surface coverage and variations in microbial reduction achieved under different locations wetted by sanitizers but with different spray exposures has not been well addressed. Surfaces that were only contacted by sanitizer via the fluid flowing down from the point of impingement fell short of achieving a 5-log reduction even when contact time was increased to 120 s. Therefore, the efficacy of sanitizer application is, in practice, likely less than what is achieved in bench-scale experiments given the reasonable possibility that individual operators will not treat all surfaces with the direct impingement of the sanitizer spray. Furthermore, research quantifying the efficacy of sanitizers may overestimate microbial reductions if reduction is only measured from coupons submerged in sanitizer or sanitizer application in sprayers but only evaluated at the point of impingement (Bae et al., 2012; Beltrame et al., 2015; Cruz & Fletcher, 2012; Gibson et al., 1999). Notably, increasing the spraying time of 120 s did increase the microbial reduction at the “Fluid-film” location to over 6-log, which may be a result of prolonged exposure to shear forces and the continuous input of sanitizer flow. However, it is unrealistic for operators to precisely control the coverage area of the impinging sanitizer flow during sanitation and increasing the application time of sanitizer spraying to this extent across all surfaces during wet cleaning.

### 3.4 Manually operated sanitizer application did not achieve a 5-log microbial reduction

Each researcher tasked with spray application used a slightly different strategy which was variably effective at reducing spot-inoculated *L. innocua* on a stainless steel surface. Spray pattern, application time, and nozzle distance from the stainless steel surface varied among researchers (Figure 2b; Table 4). Researcher 1 had the lowest microbial reduction (1.9 ± 0.5 log CFU/surface), and they also had the shortest application time (6 s) as well as the longest distance between the spray nozzle and the surface (150 cm). Researcher 2 applied sanitizer for 15 s with a nozzle distance of 45 cm achieving a reduction of 3.7 ± 0.7 log CFU/surface; Researcher 3 applied sanitizer for 13 s with a nozzle distance of 75 cm, resulting in a reduction of 2.1 ± 0.6 log CFU/surface (Table 4). Each treatment by a researcher was followed by a contact time of 120 s once spraying ceased prior to swabbing and neutralization. One-way ANOVA followed by a subsequent Tukey-Test indicated significant differences in microbial reduction among all three of the researchers (Table 4). These results represent the microbial reduction across the entire surface, though future research may investigate variability among the individual inoculation points on a single surface. Like the bench-scale results, these findings indicate that longer application time of sanitizer spray is important for increasing microbial reduction on surfaces, and perhaps more important than the contact time.

**Table 4.** Treatment parameters and outcomes for 3 researchers operating the sprayer for sanitizing. Values in a column denoted with different letters are statistically different from one another (p < 0.05; ANOVA; Tukey-HSD)

| Researcher | Spray Time (s) | Contact Time (s) | Nozzle Distance (cm) | Average Log-Reduction (SE) |
| --- | --- | --- | --- | --- |
| 1 | 6 | 120 | 150 | 1.9 (0.5)<sup>A</sup> |
| 2 | 15 | 120 | 45 | 3.7 (0.7)<sup>B</sup> |
| 3 | 13 | 120 | 75 | 2.1 (0.6)<sup>C</sup> |

**Figure 2.**
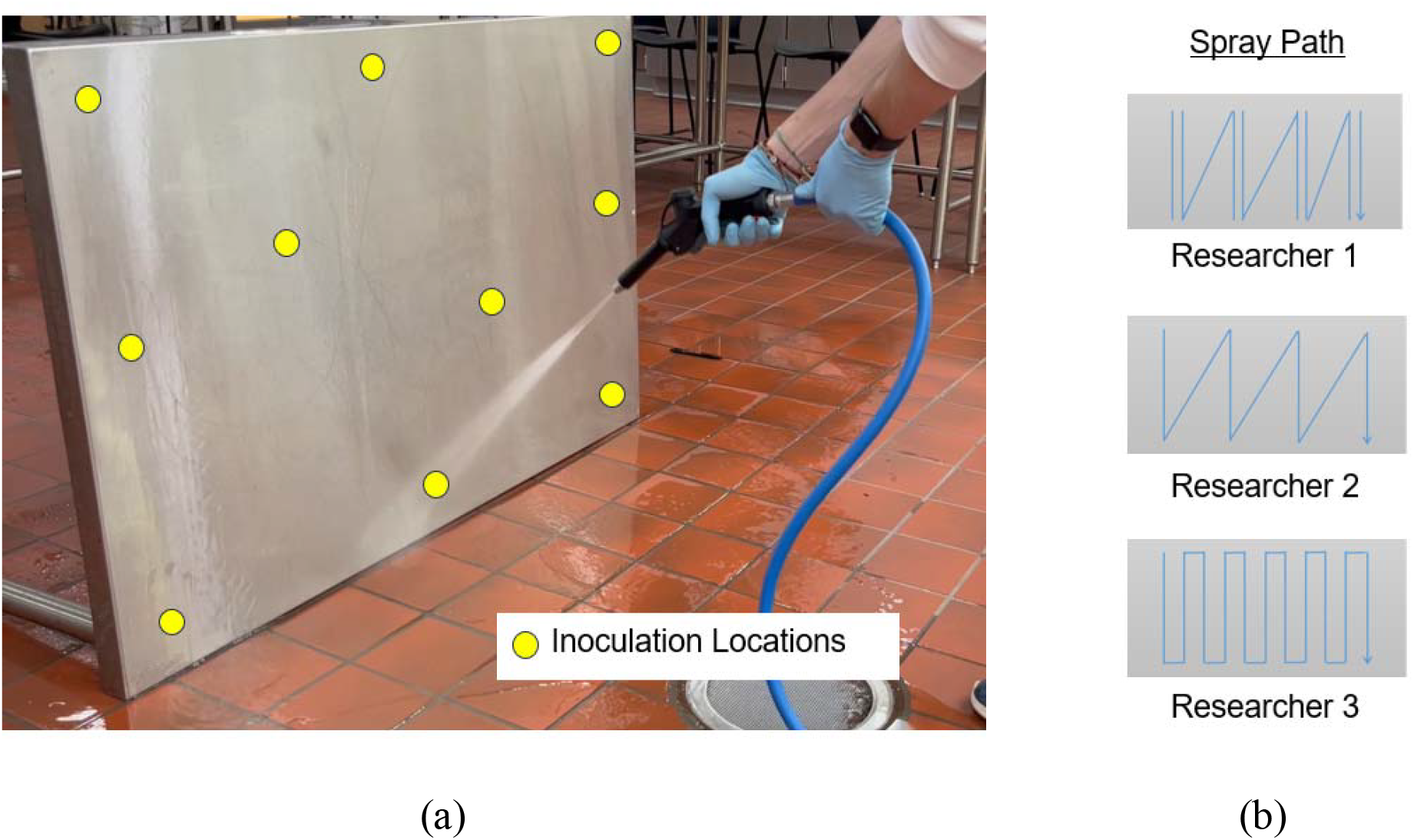
(a) Experimental set-up for pilot-scale sanitization experiment, denoting the inoculation locations (n=10) displayed in yellow. (b) Spray paths for Researcher 1, Researcher 2, and Researcher 3.

Our bench-scale experiments showed that with the correct treatment parameters both spray application and submersion in sanitizer were able to achieve a 5-log reduction on a stainless steel surface. However, in the pilot-scale experiment all three researchers failed to achieve a 5-log reduction through spray application of sanitizer (Figure 5). This was due in part to the extent that all inoculation sites on the stainless steel surfaces were treated by impinging sanitizer spray as opposed to wetted by fluid film. Outcomes were likely also influenced by other highly variable treatment factors such as nozzle distance from the surface and spray application time which change among human operators.

**Figure 5.**
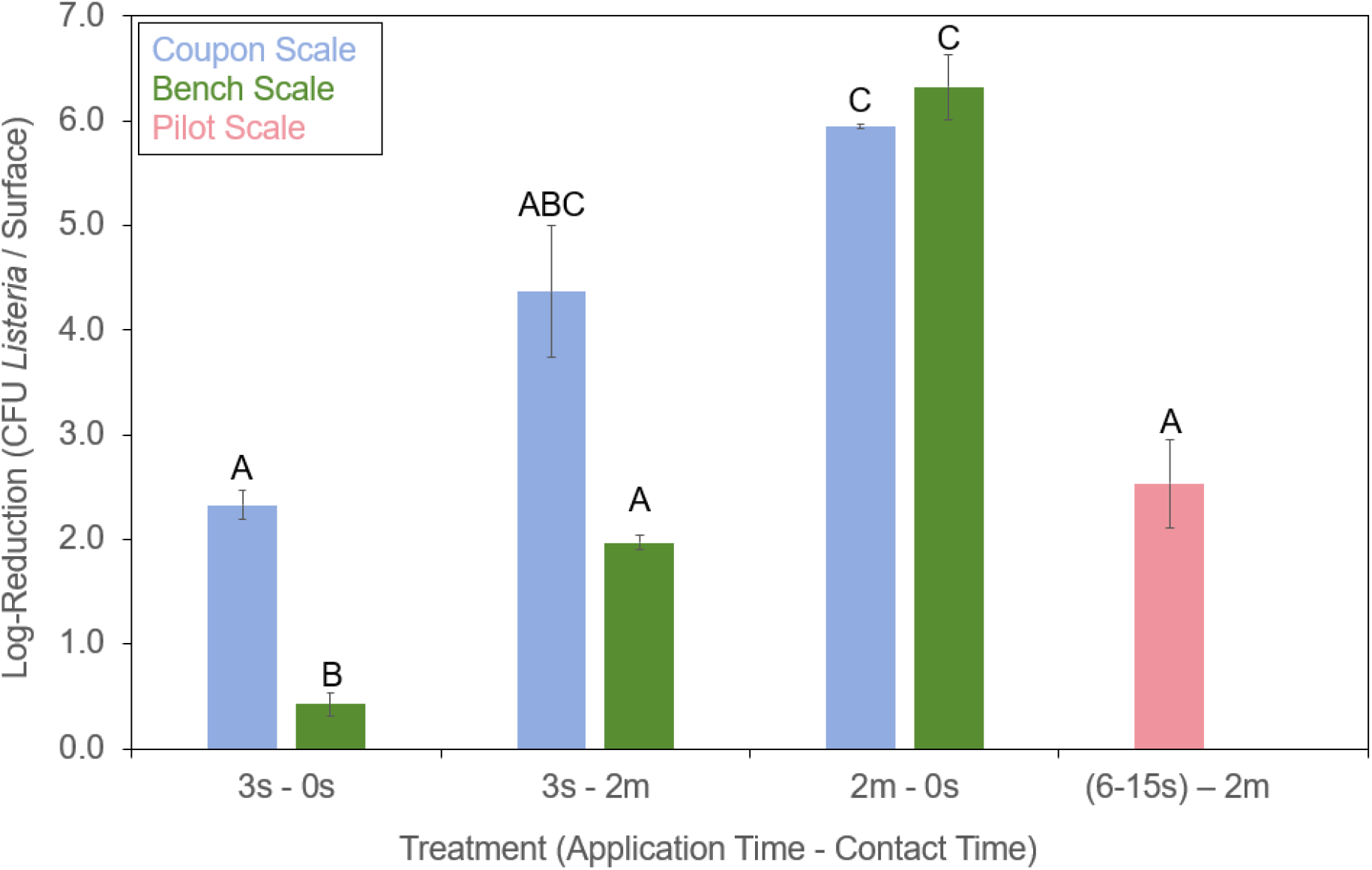
Reduction of *L. innocua* from sanitizer treatment under three conditions: coupon submersion (blue), sprayers in a bench-scale experiment (green), and researchers using sprayers at the pilot-scale (pink). Data from the “Fluid-film” location in the bench-scale experiments is presented in this figure (green). Bars denoted with different letters are statistically different from one another (p < 0.05; Welch’s-ANOVA; Games-Howell)

These results highlight how submerged coupon and bench-scale scale experiments may overestimate microbial reduction and may not be reflective of sanitation efficacy in more dynamic environments such as in pilot plants or full-scale industrial facilities. Baker et al. (2025) similarly found that application of a superheated steam sanitation treatment on bench-scale surfaces was much higher than what was achieved during pilot-scale application of the treatment.

Indeed, past studies enumerating bacterial populations in facilities before and after sanitation operations often find minimal microbial reduction especially in difficult to clean areas (Lehto et al., 2011; Nantob□Bikatui, 2020). It is possible that training interventions could result in measurable improvements in sanitation outcomes as found in past work in food establishments (Soares et al., 2013). However, some research has highlighted the limitation of training interventions to reduce post-pasteurization product recontamination (Reichler et al., 2020). Prior research has also shown that there is variability between individuals in what is considered a clean food contact surface during visual inspections (Daeschel et al., 2023), which could contribute to variations in cleaning and sanitation thoroughness among sites. Implementation of other tests to ensure sanitation operations are achieving meaningful reductions on environmental surfaces can help reduce variation in sanitation efficacy and improve hygienic conditions (Chen et al., 2022; Sogin et al., 2021).

## Conclusion

Our study elucidates the relationship between microbial reduction from sanitizing surfaces and the physical spraying parameters such as application time and method, contact time, proximity to the point of impingement, and shear stress. Shear stress varied among surface locations during sanitizer spay, which caused significant difference in *Listeria* reduction across those locations. Specifically, we noticed the greatest microbial reduction on surfaces at, and adjacent to, the impingement point where shear stress was high. Bench-scale experiments measuring the efficacy of sanitizers may overestimate sanitation outcomes compared to real world conditions where sanitizer sprayers are manually operated. The complexity of sanitizer spray effects on environmental surfaces, influenced by numerous chemical and physical variables, underscores the need for future research examining the impacts of different treatment times, spray distances, and angles on various surface materials to optimize sanitizer spray efficiency in food plant environments.

## Acknowledgements

This research was supported by the U.S. Department of Agriculture, National Institute of Food and Agriculture project 2022-67017-36289.

## Notes

### Competing Interest Statement

The authors have declared no competing interest.

